# High variability and limited host specificity characterise the microbiome of *Desmophyllum pertusum* across a cold-water coral reef

**DOI:** 10.64898/2026.10.15.759316

**Authors:** Chloé Stévenne, Christian Pruckner, Ann I. Larsson, Susanna M. Strömberg, Nadjejda Espinel-Velasco, Sofie D’hondt, Elisabeth Haring, Martin Kapun, Pedro R. Frade

**Author notes:** **Correspondence:**Chloé Stévenne, Pedro Frade.

## Abstract

Cold-water corals (CWC), like their tropical counterparts, are ecosystem engineers that create complex habitats supporting high biodiversity. Despite their broad distribution and increasing exposure to climate change and anthropogenic stressors, CWC remain comparatively understudied, particularly with respect to their associated microbiomes. Unlike tropical corals, CWC lack photosynthetic algal symbionts, suggesting that microbial associates may play a particularly important role in nutrient acquisition, biogeochemical cycling, and host resilience. Here, we investigated microbial community composition across the reef ecosystem formed by the cosmopolitan CWC *Desmophyllum pertusum* at Tisler Reef (NE Skagerrak, Kosterfjord, Norway). Using Nanopore sequencing of the full-length 16S rRNA gene, we characterised microbial communities associated with coral, seawater, substrate, and coral rubble to (i) assess reef-scale patterns, (ii) characterise variation in microbial assemblages between coral colonies and between branch and mucus compartments, as well as according to branch position, and (iii) identify putative core microbiome members and indicator taxa associated with *D. pertusum* relative to its surrounding environment. At the reef scale, microbial community structure differed among habitat compartments. Within *D. pertusum*, both colony identity and compartment (branch vs. mucus) contributed to structuring microbial assemblages, while branch position also explained an additional but limited proportion of the variation. The coral-associated microbiome contained a set of recurrent taxa, but many of these were also widespread in the surrounding environment, and few were consistently enriched in coral samples. Together, these findings indicate that the *D. pertusum* microbiome at Tisler Reef comprises recurrent microbial associates embedded within a highly variable assemblage, rather than a large and abundant core microbiome exclusive to the coral habitat. These results contribute to establishing a baseline understanding of *D. pertusum* microbiomes and provide insights into microbial community structure across cold-water coral reef ecosystems.

## 1 Introduction

Cold-water corals (CWC) are widespread across temperate, tropical and polar latitudes, forming reef structures in continental margins and seamount settings where they act as ecosystem engineers and form highly valuable habitats (Cordes et al., 2023). One of the most widespread engineer species of cold-water reefs is the coral *Desmophyllum pertusum* (formerly *Lophelia pertusa*), which has been studied across numerous locations and habitats such as Norwegian cold-water reefs (Fosså et al., 2002; Costello et al., 2005; Dorey et al., 2020), Mediterranean plateaux and submarine canyons (Taviani et al., 2004; Yakimov et al., 2006; Meistertzheim et al., 2016), Western Atlantic seamounts (Kellogg et al., 2017; Weinnig et al., 2024; Zanotti et al., 2024), or the Gulf of Mexico (Brooke and Young, 2009; Kellogg et al., 2009; Georgian et al., 2014). *D. pertusum* corals exhibit a range of morphologies and sizes, from isolated colonies in small patch reefs to massive three-dimensional structures forming large reefs and carbonate mounds built over millions of years by successive generations of colonies (Roberts et al., 2006; Sanna and Freiwald, 2021). These frameworks create structurally complex habitats that support elevated local biodiversity by providing shelter, nursery grounds, and feeding areas for a wide range of fish species (Costello et al., 2005), benthic macro- and megafauna including molluscs, arthropods, annelids, echinoderms, as well as sponges and cnidarians (Jonsson et al., 2004; Demopoulos et al., 2014).

Like their shallow-water tropical counterparts, CWC reefs are increasingly threatened by anthropogenic pressures. Human activities such as bottom trawling and deep-sea extractive industries have been shown to cause extensive damage to these habitats (Fosså et al., 2002; Ragnarsson et al., 2017), and although some regulation and conservation measures are implemented to protect them from further degradation (Huvenne et al., 2016; Otero and Marin, 2019), the ever-increasing effects of climate change on the marine environment add further pressure on these fragile ecosystems (Roberts and Cairns, 2014; Morato et al., 2020; Weinnig et al., 2020). Deep-sea biotopes remain poorly characterised and largely understudied compared to shallow tropical-water environments. Yet, the growing body of research on *D. pertusum* reefs has helped highlight the numerous regulating and provisioning ecosystem services these biotopes provide through nutrient cycling, chemosynthetic primary production, secondary production, and habitat provisioning (La Bianca et al., 2026). Beyond their role as habitat-forming species, CWC also play a vital role in biogeochemical cycling and have been recognized as hotspots for organic carbon cycling, nitrogen cycling, and benthic respiration in the deep sea (White et al., 2012; Cathalot et al., 2015; De Clippele et al., 2021). At Tisler Reef, a large reef structure in the North-East Skagerrak, the mucus-associated microbial communities of *D. pertusum* have been shown to contribute to organic matter degradation and to influence microbial activity in the surrounding water column (Wild et al., 2008, 2009).

Microbiomes of corals are integral components of the coral holobiont (Reshef et al., 2006; Bourne et al., 2016), contributing to nutrient acquisition, pathogen defence, and overall host fitness, and their study holds considerable potential for better understanding holobiont functioning and resilience (Van Oppen and Blackall, 2019; Voolstra et al., 2024). Microbiome flexibility has been proposed as a mechanism facilitating coral acclimation to changing environmental conditions on timescales far shorter than those required for host adaptation (Voolstra and Ziegler, 2020). Microbial associations may be particularly relevant for CWC, as they lack the photosynthetic Symbiodiniaceae partners that their tropical counterparts heavily rely on for nutrition. Consequently, CWC microbiomes have been hypothesised to play comparatively larger roles in nutrient acquisition and cycling (Yakimov et al., 2006; Neulinger et al., 2008; Galkiewicz et al., 2011; Kellogg et al., 2017). Despite this, CWC microbiomes remain far less studied than those of tropical corals, and our understanding of their diversity and functional roles is limited. Putative roles in biogeochemical cycling and nutrition have been investigated (Middelburg et al., 2015; Osman et al., 2023), and existing evidence indicates that CWC-associated microbial communities can be measurably affected by diet (Galand et al., 2020), as well as changes in environmental conditions, including temperature (Chapron et al., 2021; Chemel et al., 2024), and that such shifts in microbiome composition can be indicative of declines in host health (Chapron et al., 2021; Chemel et al., 2024).

Research on the microbiome of *D. pertusum* spans two decades and a range of methodological approaches of varying taxonomic resolution, from early culture-independent 16S rRNA gene clone libraries (Yakimov et al., 2006) and community fingerprinting methods such as T-RFLP, ARISA, and DGGE (Neulinger et al., 2008; Kellogg et al., 2009; Schöttner et al., 2009, 2012), through culture-based surveys (Galkiewicz et al., 2011), to amplicon sequencing on 454 pyrosequencing and Illumina platforms (Van Bleijswijk et al., 2015; Meistertzheim et al., 2016; Kellogg et al., 2017; Jensen et al., 2019; Chapron et al., 2020, 2021; Chemel et al., 2024; Zanotti et al., 2024). Across this body of work, *D. pertusum* consistently harbours microbial communities that differ from those in the surrounding environment (Yakimov et al., 2006; Neulinger et al., 2008; Schöttner et al., 2009, 2012; Van Bleijswijk et al., 2015), yet these communities show considerable variability among colonies, colour morphs, and sites (Neulinger et al., 2008; Schöttner et al., 2012; Jensen et al., 2019), and in some cases, seasons (Meistertzheim et al., 2016). The degree of host specificity also remains debated: some studies report a set of conserved, geographically widespread core taxa (Kellogg et al., 2017), or species-specific taxa associated with *D. pertusum* in CWC multi-species comparisons (Zanotti et al., 2024), whereas other studies report weak host specificity and pronounced biogeographic imprinting of *D. pertusum*’s associated microbial communities (Schöttner et al., 2012; Meistertzheim et al., 2016; Chapron et al., 2020). Despite this variability, several microbial phyla are recurrently reported as abundant across studies, including Pseudomonadota (mostly Alphaproteobacteria and Gammaproteobacteria), Actinomycetota, Bacillota, Bacteroidota, Planctomycetota, and more occasionally, Spirochaetota (Chapron et al., 2020, 2021; Chemel et al., 2024; Galkiewicz et al., 2011; Jensen et al., 2019; Kellogg et al., 2009, 2017; Meistertzheim et al., 2016; Neulinger et al., 2008, 2009; Van Bleijswijk et al., 2015; Yakimov et al., 2006; Zanotti et al., 2024; see ***Table 1*** for a comprehensive summary of taxa reported across the literature).

**Table 1.**
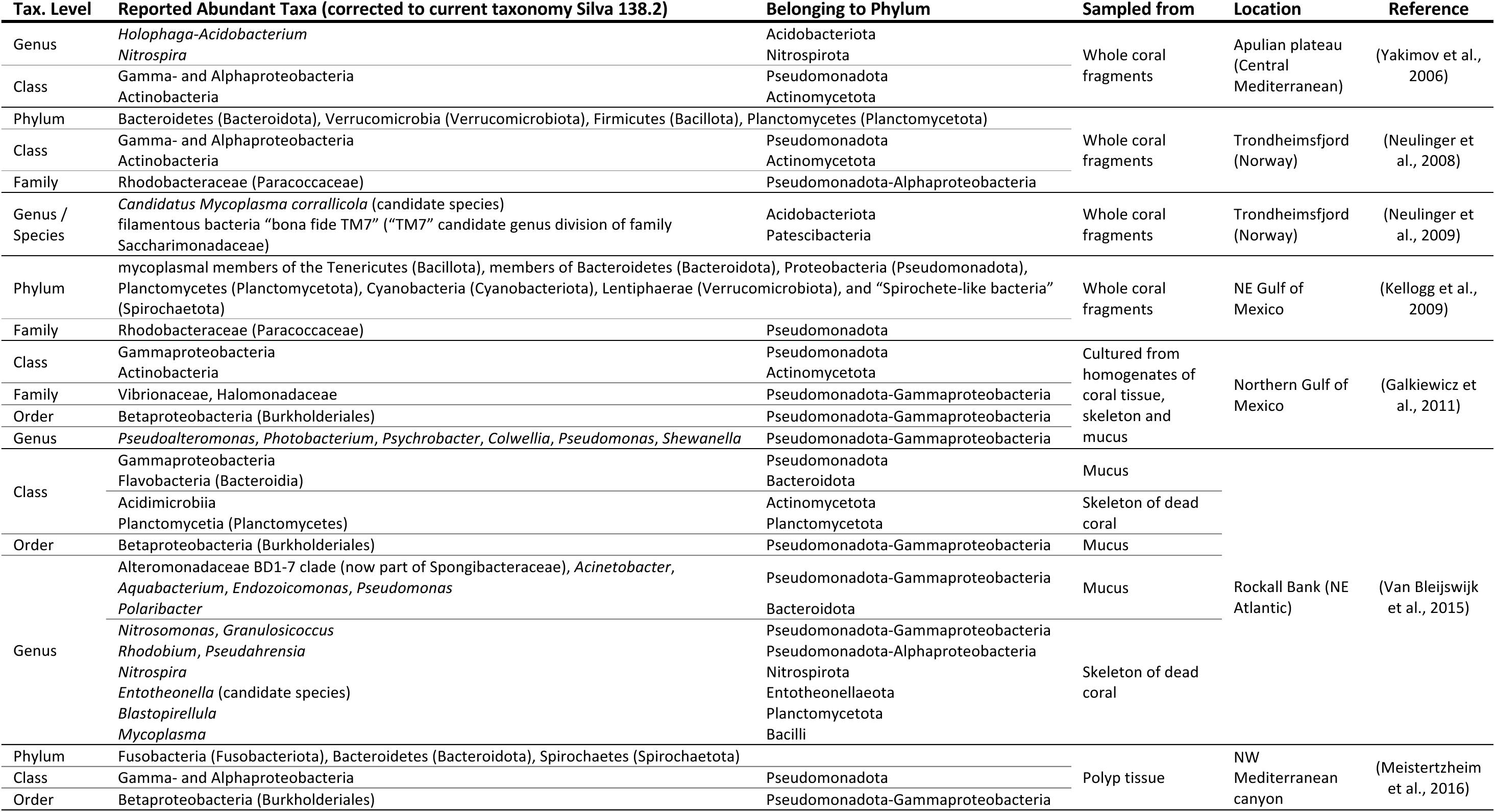
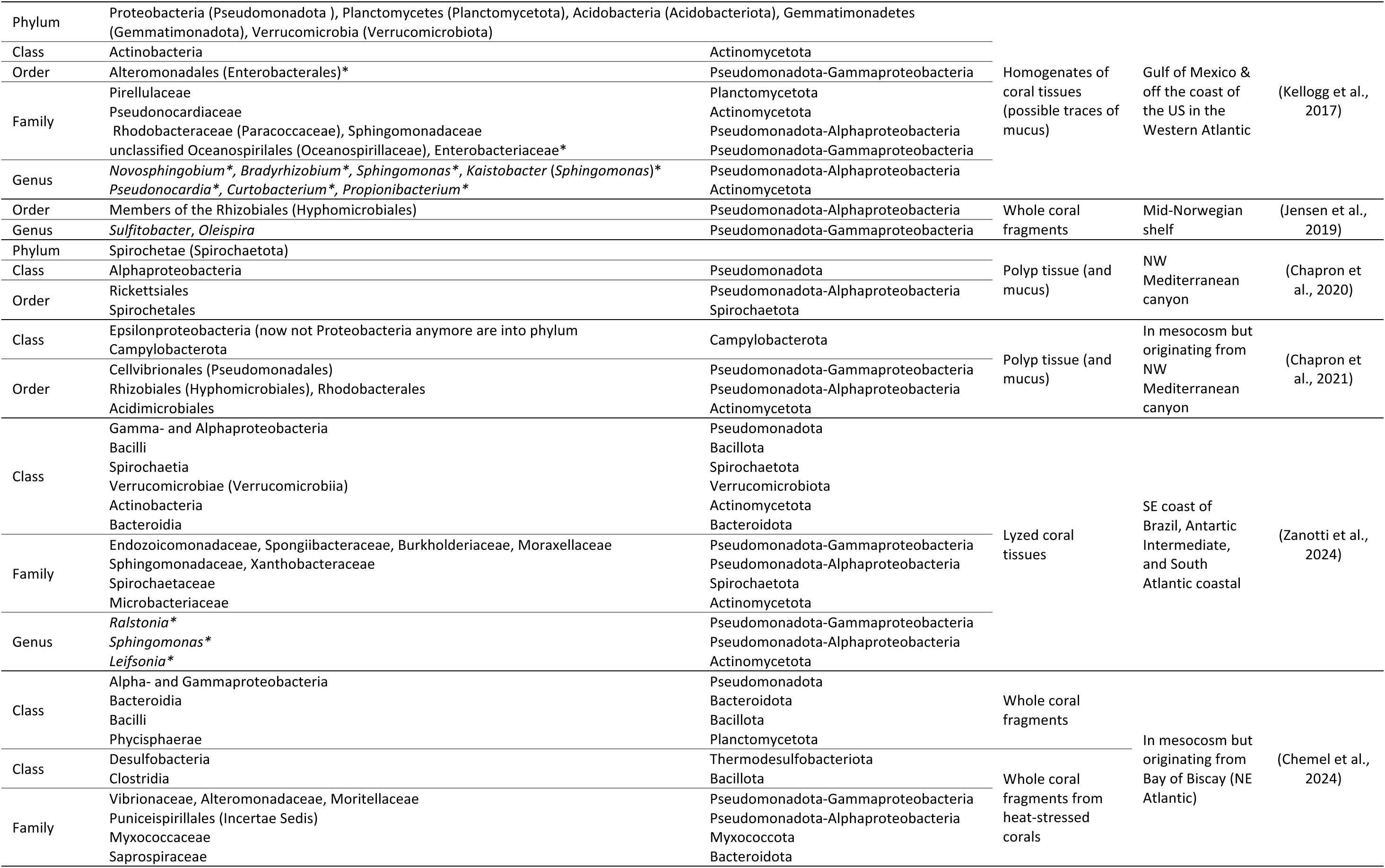
Summary of taxa reported in the literature as abundant in association with *D. pertusum.* The taxa are grouped by taxonomic level and listed as they are reported in the literature. Correction to current taxonomy according to the Silva 138.2 taxonomic database is written in parenthesis when necessary. The associated phylum membership is reported in the third column. In the case of the phylum Pseudomonadota, the class is also specified. Taxa identified as core taxa in the study are marked by an asterisk (*).

Taken together, the current literature on *D. pertusum* indicates that the coral harbours microbial communities that diverge from the surrounding environment yet are likely shaped by both host- and site-specific factors. Although some taxa have been reported as consistently dominant across several studies, clear microbial signatures for *D. pertusum* remain elusive. Earlier studies have relied on techniques offering coarse taxonomic resolution, and few have simultaneously compared the coral-associated microbial communities to those of the different habitats present on a reef, including coral branches, mucus, dead coral rubble, substrate, and water column, within a single sampling design. Consequently, the extent to which *D. pertusum* harbours microbial taxa that are consistently and specifically associated with the living coral, as opposed to taxa that are widespread across the reef environment, remains only partially resolved. Although Tisler Reef (NE Skagerrak) is one of the best studied cold-water coral reefs, only coral mucus has previously been characterised, as part of a broader four-reef comparison using ARISA fingerprinting (Schöttner et al., 2012). Here, we used full-length 16S rRNA gene sequencing on the Oxford Nanopore platform to characterise the microbial communities associated with *D. pertusum* at Tisler Reef. We compare microbial communities across coral branches (sampled according to branch position), coral mucus, dead coral rubble, reef substrate, and the overlying seawater from three colonies in order to (i) characterise reef-scale variation in microbial community composition; (ii) determine the extent to which microbial communities associated with *D. pertusum* differ between colonies, between branch and mucus compartments, and according to position on the branch; and (iii) identify putative core microbiome and indicator taxa specifically and consistently associated with the living coral host, relative to its surrounding environment.

## 2 Material and methods

### 2.1 Site description and sample collection

Sample collection took place in November 2022 at the Tisler Reef (NE Skagerrak), one of the largest structures in the Kosterfjord area (***Fig. 1A-B***). This reef forms a SE-NW oriented sill between the Tisler islands and the coast of Norway (Wisshak et al., 2005; De Clippele et al., 2018). Sampling was performed on the southeastern side of the sill using a remotely operated vehicle (ROV) V8 M500 equipped with a manipulator arm and a four-compartment sample tray (***Fig. 1C-E***). Samples were taken from three colonies from reef patches separated by less than 60 m and located between 110 and 120 m depth (Colony A: 110 m, 58°59’44.16’’N, 10°58’10.20’’E; Colony B: 114 m, 58°59’43.08’’N, 10°58’13.44’’E; Colony C: 119 m, 58°59’43.44’’N, 10°58’10.92’’E). Fragments from one healthy colony were collected at each site, along with fragments from coral rubble and adjacent substrate. The latter consisted of sediment at Colony A and rock or shell material at Colonies B and C). The samples were brought onto the research vessel Nereus of the Tjärnö Marine Laboratory (University of Gothenburg), where they were handled with sterile tools. Each coral sample consisted of a large fragment comprising several branches. Mucus samples were collected from each colony using sterile cotton swabs gently rolled over the coral surface. The coral fragments were then divided into smaller branch fragments (typically 1-2 cm long), which were sorted into three categories according to their position on the collected fragment: branch tips (containing a polyp), middle sections (interbranch regions without polyps), and basal sections (thicker regions where several branches joined). Seawater was sampled at each site using niskin bottles and 5 L were filtered onto a 25 mm polycarbonate filter with pore size 0.2 µm (Whatman) or on a Sterivex^TM^ filter (Millipore) using a peristaltic pump. Coral branches subsamples, mucus cotton swabs, substrate samples, and filters were immerged in DESS solution (0.25M Na_2_EDTA, 20% DMSO, NaCl saturated, pH 8.0) or DNA/RNA Shield (Zymo Research, USA) and preserved at 4°C until they were brought to the laboratory facilities and placed at -20°C. Across the three sites, a total of 47 subsamples from healthy coral branches were collected and retained for analysis, alongside 17 mucus swabs, 11 samples from coral rubble branches, 7 substrate samples, and 9 seawater samples (***Supplementary Table S1***).

**Fig. 1.**
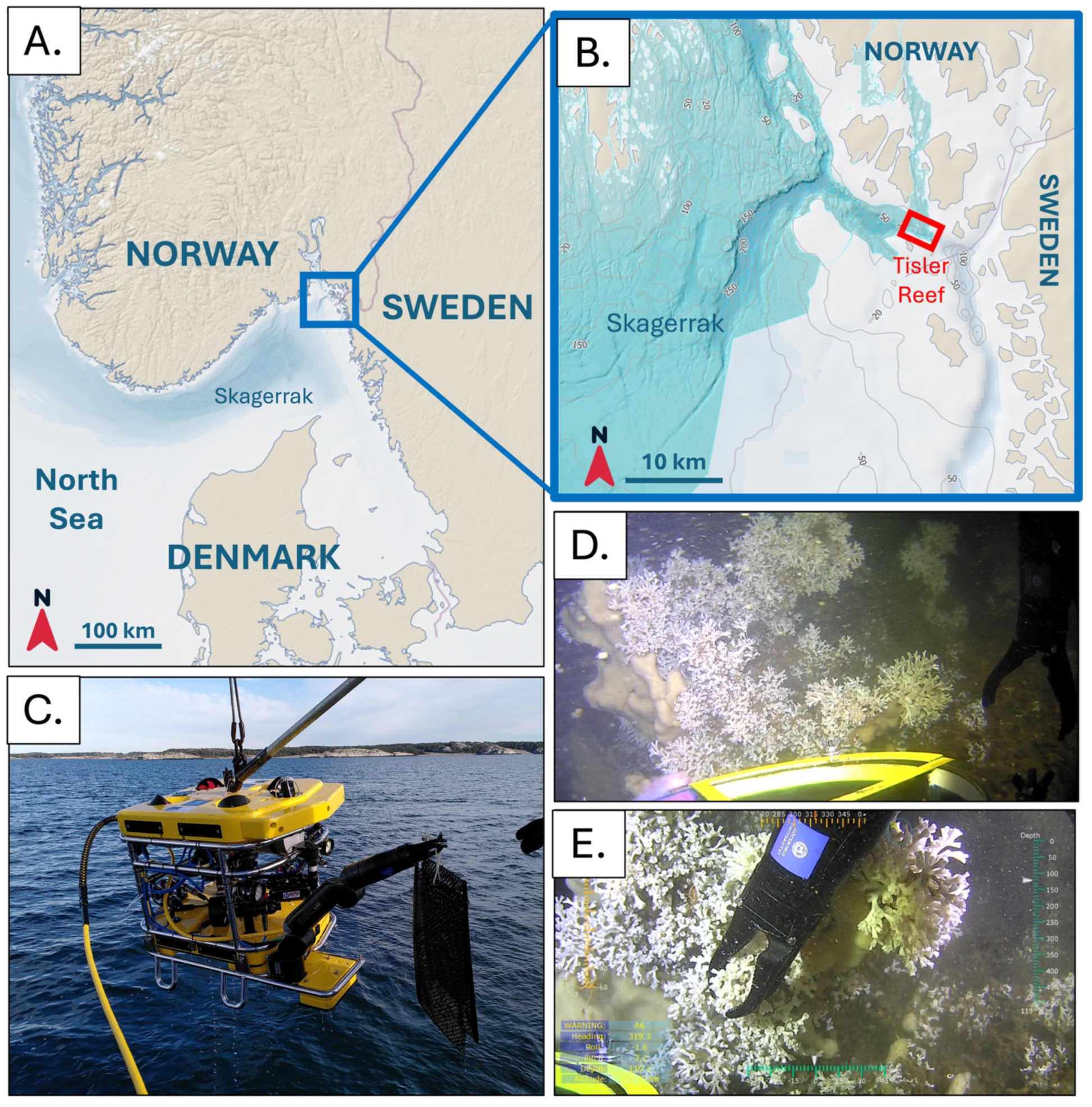
Sampling campaign on Tisler Reef. **A-B.** Maps showing the location of Tisler Reef in the northeastern Skagerrak, between Norway and Sweden. The reef forms a SE-NW oriented sill between the Tisler islands and the Norwegian coast.**C.** ROV V8 M500 used for sampling at depth. **D.** Approach of colony C during the 3^rd^ sampling dive. **E.** Sampling of healthy *D. pertusum* branches from colony C using the manipulator arm of the ROV.

### 2.2 DNA extraction, PCR amplification and long-read sequencing

Coral branch samples were rinsed with 500 µL sterile water to remove loosely associated microbes prior to DNA extraction. All DNA extractions were performed using the ZymoBIOMICS DNA Miniprep Kit (Zymo Research, USA) following the manufacturer’s instructions, though adapting the initial step with the addition of 20 µL Proteinase-K (20 mg mL^−1^) and 42 µL 20% SDS for a 30 min digestion at 60°C. The full length 16S rRNA gene was amplified using forward primer 27F_BCtail-FW 5’-TTTCTGTTGGTGCTGATATTGC_AGAGTTTGATCMTGGCTCAG-3’ and reverse primer 1492R_BCtail-RV 5’-ACTTGCCTGTCGCTCTATCTTC_CGGTTACCTTGTTACGACTT-‘3 following the protocol established by van der Loos et al. (2021). Briefly, PCR reactions including negative controls were performed using the Phire Tissue direct PCR Master Mix (Thermo Scientific) in 15 µL reactions and consisted of an initial denaturation at 98°C for 3 min, followed by 30 cycles of 98°C for 8 sec, 60°C for 8 sec, 72°C for 30 sec, and a final extension at 72°C for 3 min. Library preparation consisted in amplicon barcoding using the Oxford Nanopore Barcoding Expansion 1-96 kit (EXP-PBC096) with an initial denaturation at 95°C for 3 min, followed by 13 cycles of 95°C for 15 sec, 62°C for 15 sec, 65°C for 40 sec, and a final extension at 65°C for 2 min. PCR purification using Agencourt AMPure XP Beads and the ligation sequencing kit SQK-LSK114 (Oxford Nanopore Technologies) was used for the final library preparation step prior to sequencing on a MinION with an R10.4.1 flow cell for 72h (van der Loos et al., 2021). Dorado (v7.3.11, Oxford Nanopore Technologies) was used for super-accurate base-calling. The minimum data quality threshold was set to Q10.

### 2.3 Read processing, taxonomic assignment and statistical analyses

Data quality and read length were assessed with NanoPlot (v1.32.1) before filtering was applied based on length (1,300 bp ≤ read length ≤ 1,800 bp) and quality (minimum Phred average quality ≥ 12) with Chopper (v0.9.0)(De Coster and Rademakers, 2023). The resulting high-quality reads were then taxonomically classified using Emu (v3.4.6), which employs an expectation-maximization algorithm to generate taxonomic abundance profiles from full-length 16S rRNA gene reads (Curry et al., 2022). The SILVA v138.2 16S rRNA gene database (Quast et al., 2013), trained for DADA2 (Callahan et al., 2016), was reformatted into FASTA and mapping files to construct a custom database compatible with Emu (Curry et al., 2022). This ensured the use of the most up-to-date taxonomic nomenclature as implemented in the latest SILVA database release (SILVA, 2024).

The resulting taxonomic abundance profile table obtained from Emu was uploaded in R (v4.3.1)(R Core Team, 2018). Each entry in the resulting abundance table, corresponding to a taxonomic assignment in the reference database, was considered as a taxonomic feature (TF) or taxon in the downstream analyses. Taxonomic features assigned to eukaryotes, mitochondrial, or chloroplast DNA were removed from the dataset. Negative controls introduced during the first PCR were used to detect and remove potential contaminants, which were identified using the *decontam* package (v1.20.0)(Davis et al., 2018), applying the prevalence method with a threshold of 0.2. Finally, the resulting decontaminated abundance table was rarefied to an even sequencing depth of 28,451 reads. All subsequent analyses and statistics were conducted in R using the *phyloseq* (McMurdie and Holmes, 2013) and *vegan* (Oksanen et al., 2015) packages, and graphical representations were generated using *ggplot2* (Wickham, 2009).

Visualization of the microbial communities were performed on rarefied relative abundance data to assess taxonomic composition. Relative abundance values are reported as mean ± standard deviation (SD). Log_10_ transformed data were used for Principal Coordinates Analysis (PCoA) based on Bray-Curtis dissimilarities to illustrate beta-diversity of microbial communities across the different sample types. PERMDISP and PERMANOVA (9,999 permutations) were used to test homogeneity of multivariate dispersions and to evaluate the individual and combined effects of sample type and colony site on microbial communities at the reef-scale. At the coral-scale, they were likewise used to test the effects of colony and sample type (branch vs. mucus), and, for branch samples only, of colony and branch position (tip, middle, basal). Colony (n = 3) was treated as a fixed factor, and its effect was tested with unrestricted permutations. To account for non-independence among samples from the same colony, sample type and branch position were also tested in additive models with permutations restricted within colonies. As branch samples were collected randomly from multi-branch colony fragments and assigned a posteriori to position categories, non-independence among samples based on position could not be accounted for. Alpha-diversity was measured as observed richness, evenness and as the Shannon index calculated from the rarefied raw sequence data. Diversity estimates were calculated over 100 iterations and averaged across all trials. The effects of sample type and colony site on alpha-diversity were assessed with a two-way ANOVA followed by a post-hoc Tukey HSD test. Variables that did not meet normality assumptions were measured using the non-parametric Kruskal-Wallis rank-sum test followed by a post-hoc Dunn test. Because substrate material differed among sampling locations, substrate samples were not considered as distinct substrate types in the statistical analyses. Core microbiome taxa associated with coral branch samples across the three colonies were defined as the taxa with a minimum prevalence of 90% across all samples (n=47). The prevalence of each taxon was assessed and visualized across a range of detection thresholds (relative abundances from 0 to 0.3%) using the *plot_core* function from the R microbiome package (v1.22.0). Indicator species analysis was conducted using the *multipatt* function with 999 permutations of the *indicspecies* package in R (De Cáceres and Legendre, 2009). Indicator taxa were filtered based on the thresholds p-value ≤ 0.01 with minimum specificity (component A) and minimum fidelity (component B) set to 0.5.

## 3 Results

### 3.1 Overview of sequencing and 16S rRNA gene data processing

A total of 12,161,473 reads passed quality filtering (Q > 10) across 109 samples, which included all primary samples, technical duplicates for low-DNA-yield samples, one mock community, and negative controls. This corresponded to an average of 111,573 reads per sample. After taxonomic assignment, additional filtering on taxonomy, and contaminant removal, 8,155,219 classified reads remained across 105 samples, for an average of 77,548 reads per sample. Rarefaction to 28,451 reads per sample resulted in the exclusion of 6 samples, only 2 of which were unique coral branch samples (one from colony A, and one from colony B) without duplicates. The other 4 excluded samples were technical replicates, with their respective paired samples retained in the dataset.

### 3.2 Habitat-scale variations of microbial communities

Microbial communities across the reef habitat including coral branches and mucus, the substrate, coral rubble, and surrounding seawater were dominated by bacterial families belonging to the phylum Pseudomonadota (formerly Proteobacteria) with Paracoccaceae (Alphaproteobacteria) being the most abundant across all sample types, especially in seawater (11.1 ± 8.4%) (***Fig. 2A***). Leptospiraceae (Spirochaetota) were particularly dominant in microbial communities associated with

**Fig. 2.**
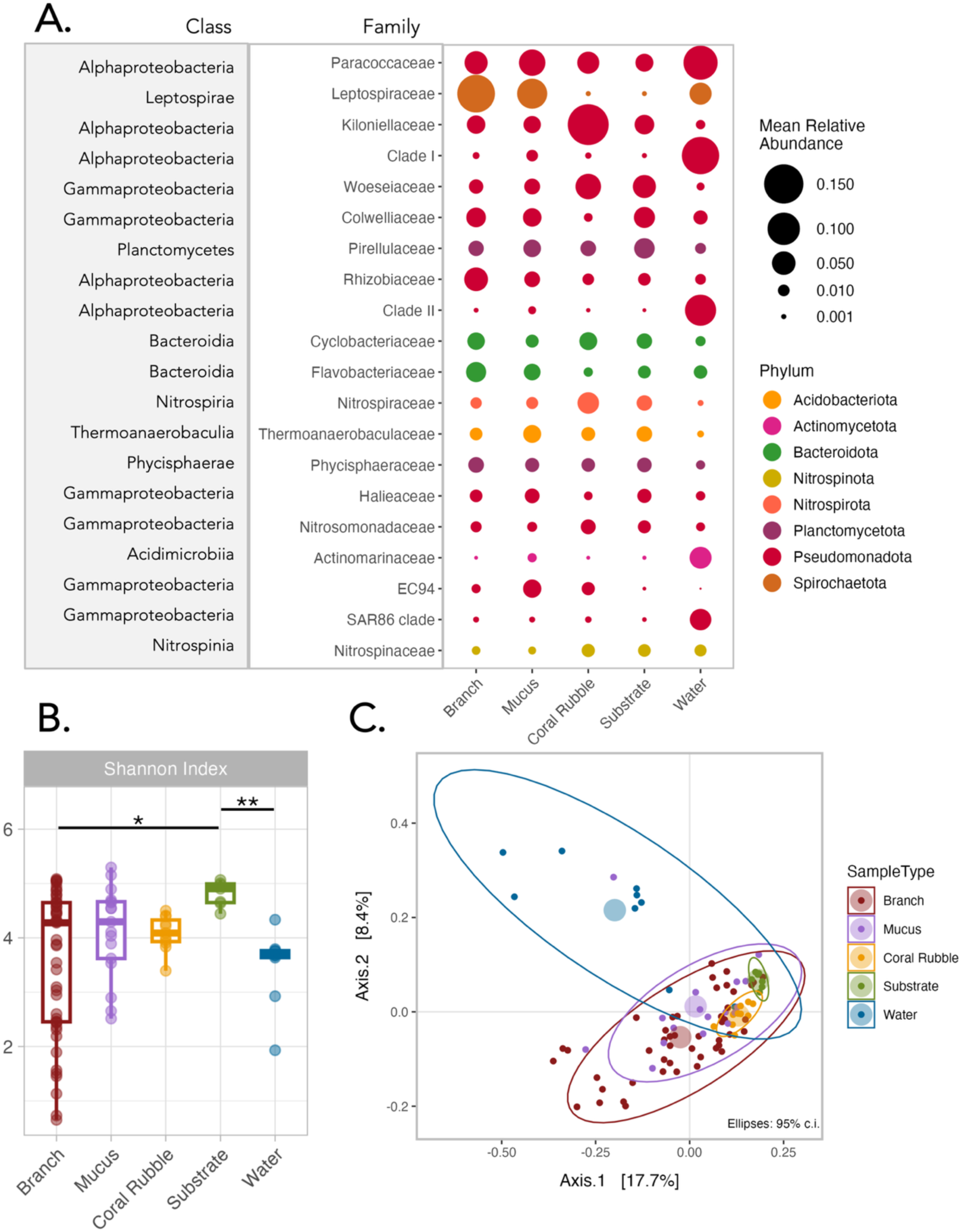
Habitat-scale variation in microbial communities associated with *D. pertusum*, surrounding substrates, and seawater. **A.** Relative abundance of the 20 most abundant prokaryotic families across all sample types, normalized by group sample size. Point size represents the mean relative abundance of each family across replicates within each sample group: coral branches (n = 47), coral mucus (n = 17), coral rubble (n = 11), substrate (n = 7), and seawater (n = 9). **B.** Alpha diversity across sample types. Shannon index calculated from rarefied read counts. Each boxplot summarizes the distribution of replicates within each sample group. **C.** Beta diversity of microbial communities by sample type. PCoA based on Bray-Curtis dissimilarities of log_10_-transformed data. Samples are coloured by sample type. Ellipses represent 95% confidence interval, and coloured disks indicate group centroids.

*D. pertusum*, both in coral branch (13.7 ± 26.9%) and mucus (8.5 ± 18.1%) samples, while their relative abundance was markedly lower in substrate and coral rubble samples (< 0.01%). In contrast, Kiloniellaceae (Alphaproteobacteria), the third most abundant family across all samples, dominated coral rubble microbial communities (16.3 ± 6.7%). Seawater communities exhibited the most distinct composition of dominant families compared to *D. pertusum*- and substrate-associated microbial communities. Families such as Pelagibacterales Clade I and Clade II (Alphaproteobacteria), Actinomarinaceae (Actinomycetota), and the SAR86 clade (Gammaproteobacteria) were more abundant in seawater, whereas groups like Kiloniellaceae (Alphaproteobacteria), Woeseiaceae (Gammaproteobacteria), Nitropiraceae (Nitrospiria), and EC94 (Gammaproteobacteria) were underrepresented relative to the other sample types.

Alpha diversity patterns revealed high variability in microbial assemblages from coral branches, mucus, and seawater, while diversity was more consistent among coral rubble and substrate samples (***Fig. 2B***). Alpha-diversity (Shannon index) was significantly influenced by sample type (Kruskal-Wallis, p < 0.01) with substrate communities showing higher diversity than those associated with coral branches (Dunn’s test, p < 0.05) and seawater (Dunn’s test, p < 0.01).

Beta-diversity (PCoA on Bray-Curtis dissimilarities; ***Fig. 2C***) showed tight clustering of substrate and coral rubble samples, and greater dispersion in coral branch, mucus, and seawater samples. This pattern was supported by a significant effect of sample type on group dispersion (PERMDISP, p = 0.0001). Dispersions in substrate and coral rubble samples differed significantly from those in the other three sample types (PERMUTEST, p < 0.05), while no significant differences in dispersion were found between coral branch, mucus, and seawater samples, or between substrate and coral rubble samples (PERMUTEST, p > 0.05). Sample type was the main driver of community variation (PERMANOVA, R^2^ = 0.16, p = 0.0001), although sampling location (coral colony for coral samples, reef patch for substrate and seawater) was also significant (PERMANOVA, R^2^ = 0.03, p = 0.0141) but not the interaction of the two variables (PERMANOVA, R^2^ = 0.07, p = 0.7398). Pairwise comparisons across all sampling locations indicated that microbial communities differed significantly between all sample types (pairwise PERMANOVA, p < 0.05).

### 3.3 *Desmophyllum pertusum*-associated microbial communities

Leptospiraceae (Spirochaetota) was the most dominant prokaryotic family in *D. pertusum* branch and mucus microbial communities across the three colonies. However, it was the most abundant family only in branch samples from colony A (8.21 ± 17.05%) and in both branch (28.83 ± 37.40%) and mucus (20.09 ± 25.97%) samples from colony C, while its relative abundance was lower in colony B (1.73 ± 4.64% in branch and 0.17 ± 0.19% in mucus) (***Fig. 3A***). The dominance of Leptospiraceae in colonies A and C was not consistent across all samples: in many, it was not the most abundant taxa, with relative abundances below 1%. In contrast, some samples from both colonies showed a strong dominance of the family, certain accounting for over 40% of the microbial community and reaching up to 92% in one case (***Supplementary Figure S2***). This pattern was especially pronounced in tip branch samples from colony C, where 5 out of 7 were dominated by Leptospiraceae, making it the only sample subgroup in which the family consistently dominated the majority of samples.

**Fig. 3.**
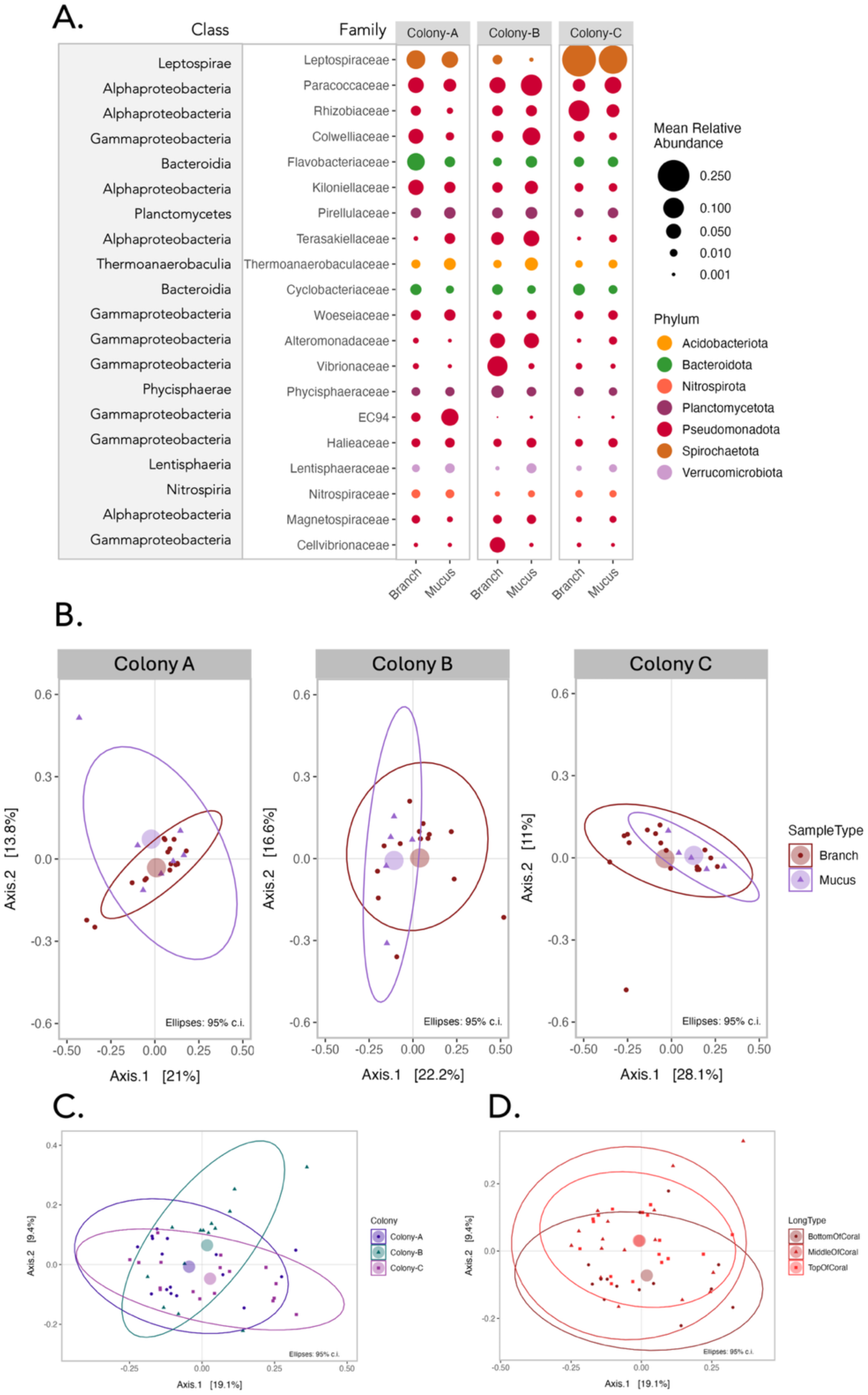
Microbial communities associated with coral branches and mucus across three colonies of *D. pertusum*. **A.** Relative abundance of the 20 most abundant prokaryotic families across all coral samples (branches and mucus), normalized by group sample size. Point size represents the mean relative abundance of each family across replicates within each sample group: coral branches (n = 47), coral mucus (n = 17). **B.** Beta diversity of microbial communities associated with coral branches and coral mucus from three different colonies. Samples are coloured by sample type and the plots are parsed according to colony. **C-D.** Beta diversity of microbial communities associated with coral branches, coloured by colony (**C**) and by position on the branch (**D**). PCoA based on Bray-Curtis dissimilarities of log_10_-transformed data. Ellipses represent 95% confidence interval, and coloured disks indicate group centroids.

Paracoccaceae (Alphaproteobacteria) were more consistently dominant in all samples from all three colonies with relative abundances ranging from 3.32 ± 3.16% in branch samples from colony C to 10.71 ± 6.98% in mucus from colony B. Rhizobiaceae (Alphaproteobacteria) were particularly dominant in some branch samples from colony C (10.40 ± 22.78%), with lower mean relative abundances in branches from colonies A (1.94 ± 1.72%) and B (2.25 ± 2.61%). Both Alteromonadaceae and Vibrionaceae (Gammaproteobacteria) were more represented in association with colony B branches than with colony A or C. These patterns were due to some branch samples with an overrepresentation of these taxa with over 50% relative abundance (***Supplementary Figure S2***). While the Gammaproteobacteria EC94 was largely absent from most branch and mucus samples from colony B and C, and from some samples from colony A, it was highly represented in a couple of branch samples (15.58% and 7.57%) and one mucus sample (46.46%) from colony A. Finally, Cellvibrionaceae (Gammaproteobacteria), although represented across most branch and mucus samples from all colonies, were particularly dominant in three branch samples from colony B at 12.13, 21.64 and 37.74% of relative abundance.

Comparing microbial communities associated with coral branches and mucus across the three colonies (***Fig. 3B***) revealed a significant effect of sample type (PERMANOVA, unrestricted: R^2^ = 0.025, p = 0.037, with permutations restricted within colonies, R^2^ = 0.024, p = 0.048) and of colony (PERMANOVA, R^2^ = 0.053, p = 0.004) on community composition. Colony was the stronger driver of the two, explaining 5.3% of the variation. Pairwise comparisons showed that Colony B was compositionally distinct from both Colony A (R^2^=0.045, p = 0.006) and Colony C (R^2^ = 0.056, p = 0.002), while Colony A and Colony C did not differ significantly from one another (R^2^ = 0.025, p = 0.244). The interaction term between sample type and colony was non-significant (PERMANOVA, R^2^ = 0.034, p = 0.232). Homogeneity of multivariate dispersion was confirmed for all groups (PERMDISP, p > 0.05), supporting the interpretation of PERMANOVA results as reflecting differences in group centroids rather than dispersion. Overall, 89% of the variation remained unexplained by the model, reflecting high within-group heterogeneity.

Microbial communities associated with coral branches were further examined according to sampling position (tip, middle, and bottom) to assess whether position along the branch influenced community composition. PERMANOVA revealed significant effects of both position (PERMANOVA, unrestricted: R^2^ = 0.064, p = 0.019, with permutations restricted within colonies, R^2^ = 0.063, p = 0.025) and colony (R^2^ = 0.076, p = 0.004), with colony remaining the stronger driver of compositional differences (***Fig. 3C***). The interaction term was non-significant (R^2^ = 0.079, p = 0.579), and 78.2% of the variation remained unexplained by the model. Pairwise comparisons showed that tip and bottom communities differed significantly from one another (R^2^ = 0.067, p = 0.009), while middle-position communities were not distinguishable from either tip (R^2^ = 0.032, p = 0.355) or bottom (R^2^ = 0.049, p = 0.083) (***Fig. 3D***). Homogeneity of multivariate dispersion was confirmed for all groups (PERMDISP, p > 0.05), supporting the interpretation of PERMANOVA results as reflecting differences in group centroids rather than dispersion.

Coral branch microbial assemblages were analysed to identify core taxa defined as taxonomic features with a minimum prevalence of 90% across all samples. A total of 42 TFs spanning nine phyla were picked out (***Fig. 4A***). Colony A and Colony C presented a similar prevalence profile of these taxa, while the profile in Colony B differed with overall lower prevalence of the identified core TFs. The majority of the identified core taxa belonged to the phylum Pseudomonadota (n=26), followed by Planctomycetota (n=6), while the other phyla were represented by one or two taxa. Nine core TFs were present in 100% of the samples across the three colonies. However, these taxa were also highly prevalent in environmental samples (100% prevalence in both substrate and coral rubble samples and minimum 66.7% prevalence in water samples). They also had relatively low abundances in coral branch samples ranging from 1.63 ± 1.27 % for the most abundant “TF-6 *Woeseia* sp.” to 0.25 ± 0.34 % for “TF-47 Clade Ia”, a Pelagibacterales, which was dominant in water microbial communities (11.54 ± 9.26 %). Across the 42 TFs identified as core taxa, five had higher mean relative abundances in coral branches than in any other sample type (TF-18 Haliangiaceae, TF-20 *Pelagibius* sp., TF-7 Rhizobiaceae, TF-41 Kangiellaceae, and TF-4 *Pseudahrensia* sp) and two were only more abundant in mucus samples (TF-13 *Roseovarius* sp. and TF-92 Legionellaceae).

**Fig. 4.**
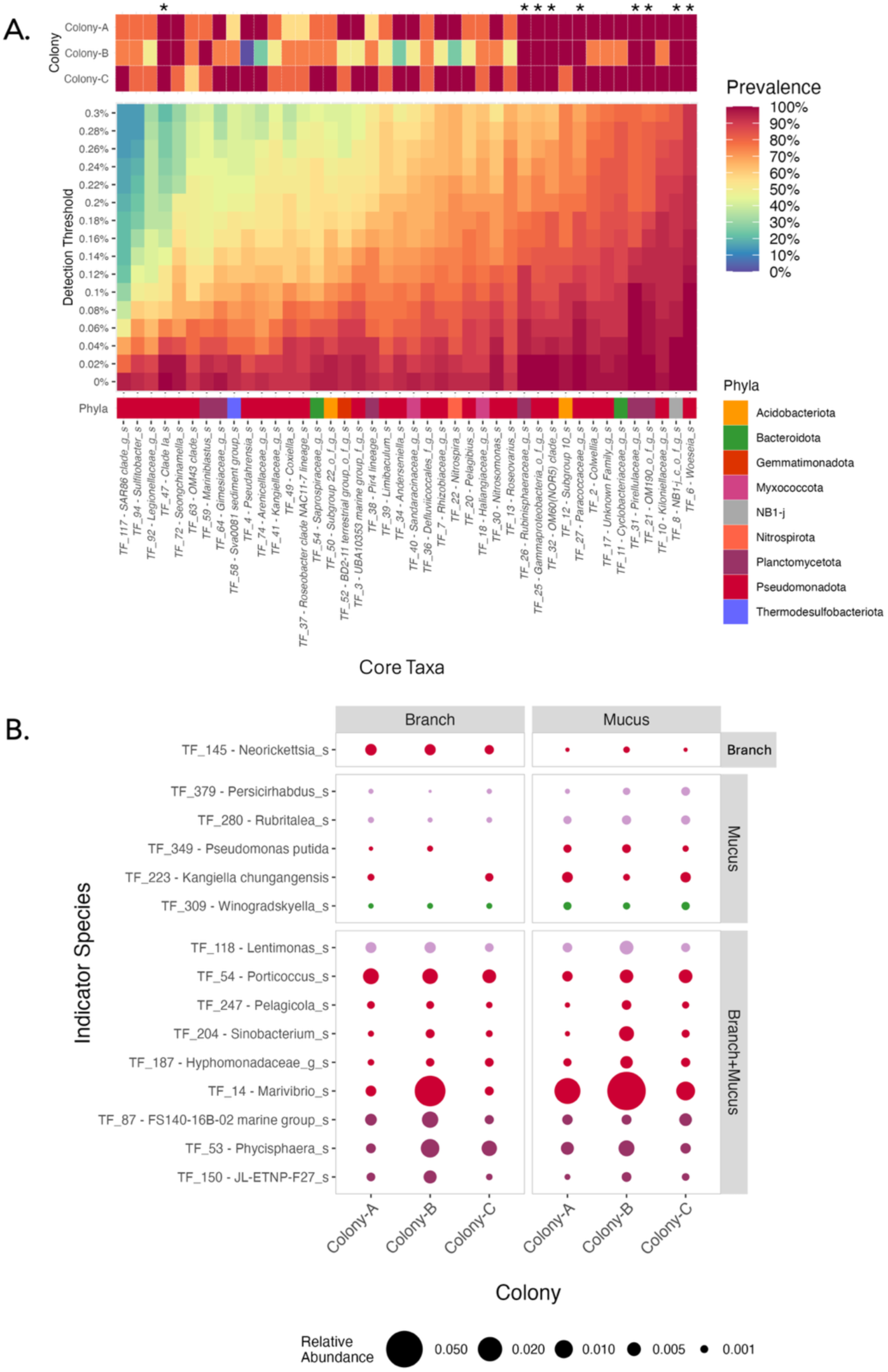
Core and indicator microbial taxa associated with *D. pertusum* across three colonies. **A.** Heatmap showing the prevalence (colour scale) of core microbial taxa across a gradient of detection thresholds (y-axis). The upper heatmap displays prevalence within each colony at a detection threshold of 0; asterisks denote the nine taxa present in 100% of samples across all colonies. The larger heatmap shows prevalence across all colonies under increasing detection thresholds. The coloured bar below indicates taxonomic affiliation at the phylum level. **B.** Relative abundance of the 15 TFs identified as indicator taxa for coral-associated samples, grouped into: branch-specific indicators, mucus-specific indicators, and shared indicators (branch and mucus). Point size represents the mean relative abundance across replicates within each sample type (branches: *n* = 47; mucus: *n* = 17), and colours indicate phylum-level classification. Taxa are defined by TF number, corresponding to their rank in relative abundance across all coral samples, followed by their lowest available taxonomic classification. Unresolved taxonomic ranks are indicated by suffixes: “_c” (class), “_o” (order), “_f” (family), “_g” (genus), and “_s” (species).

A complementary analysis was conducted on coral branch and mucus samples to identify indicator taxa defined by their specificity (the degree to which a taxon is restricted to branch and / or mucus samples) and fidelity (the frequency of occurrence within these groups). These parameters were contrasted against environmental samples and their thresholds set to 50%. This analysis picked out 15 indicator taxa: one was indicator of coral branches (TF-145 *Neorickettsia* sp. from the Anaplasmataceae family), five were indicator of coral mucus (two Verrucomicrobiota, two Pseudomonadota, and one Bacteroidota), and nine were indicator for both branches and mucus samples (one Verrucomicrobiota, five Pseudomonadota, and three Planctomycetota). Specificity values (component A) ranged between 73.1% and 97.7%, whereas fidelity values (component B) were lower, ranging between 53.1% to 89.1% (***Supplementary Table S3***). Although these taxa were selected for their combined fidelity and frequency across all coral samples, they were among lower abundance taxa with the most abundant, TF-14, at a mean relative abundance of 1.41 ± 7.16 %. Out of the 15 identified indicator taxa, five had higher mean relative abundances in coral samples than in any other sample type (TF-14, TF-53, TF-150, TF-280, and TF-379). Only TF-54 *Porticoccus* sp. was both identified as a core and indicator taxa.

## 4 Discussion

*Desmophyllum pertusum* is a widely distributed reef-forming cold-water coral, yet the extent to which it harbours a characteristic and taxonomically distinctive microbiome remains unresolved. Here, we used full-length 16S rRNA gene sequencing to characterise microbial communities associated with coral branches and mucus, dead coral rubble, substrate, and seawater at the shallow Tisler Reef (110-120 m).

### 4.1 Diversity patterns of microbial assemblages

At the reef scale, microbial assemblages differed among coral branches, mucus, rubble, substrate, and seawater compartments. Substrate-associated communities exhibited the highest Shannon diversity and the lowest among-sample compositional dispersion (***Fig. 2B-C***), indicating assemblages that were both diverse and comparatively consistent across samples. Coral rubble communities also clustered tightly, whereas seawater and coral-associated communities showed substantially greater among-sample variability. This uneven variability means the overall difference detected among habitats should be interpreted with some caution, as part of it may stem from some sampling groups simply being more heterogeneous than others, rather than reflecting a true separation between habitat types. Nevertheless, the contrasting taxonomic profiles (***Fig. 2A***) and ordination patterns (***Fig. 2B***), particularly the clear separation of seawater from benthic and coral-associated samples, are consistent with habitat-related structuring of microbial communities across the reef. These patterns broadly align with previous studies reporting habitat-specific structuring of microbial communities across seawater, sediment, and coral compartments on cold-water coral reefs, including *D. pertusum* reefs (Yakimov et al., 2006; Neulinger et al., 2008; Schöttner et al., 2009, 2012; Van Bleijswijk et al., 2015). Notably, Schöttner et al. (2012) sampled several reefs along the Norwegian continental margin, including Tisler Reef. Consistent with our results, they reported that sediment-associated assemblages combined high alpha-diversity with comparatively low beta-diversity dispersion relative to seawater and coral-associated communities (Schöttner et al., 2012). Some of the variability observed among our seawater samples may reflect differences in sampling position, as the precise distance from the seabed and coral colonies could not be fully standardised during water sampling. Microbial assemblages above CWC reefs are known to vary with height above the seabed and proximity to coral structures (Van Bleijswijk et al., 2015), making seawater samples particularly sensitive to fine-scale spatial variation.

The comparatively low dispersion of substrate and rubble communities (***Fig. 2C***) may reflect the availability of persistent surfaces for microbial biofilm development. However, the substrate samples comprised different physical materials across sampling locations, with sediment sampled at Colony A and rock or shell material at Colonies B and C. Therefore, the relatively consistent microbial signatures observed among substrate samples should be interpreted cautiously, as differences in substrate material may influence biofilm composition (Dang and Lovell, 2016) and cannot be separated from sampling location in the present dataset. The clustering of these samples may thus reflect common features of substrate biofilm-associated habitats, although the physical properties and complexity of the underlying substrate may further influence bacterial communities which could not be disentangled here. Such biofilms can support taxonomically diverse communities and generally host greater bacterial diversity than the surrounding seawater (Zhang et al., 2019; Hadfield et al., 2025). Despite their proximity and exposure to similar reef conditions, coral rubble and substrate samples diverged, suggesting that dead coral skeleton provides a distinct microbial habitat with the potential to promote overall microbial diversity at the reef scale. Microbial communities associated with coral rubble have seldom been investigated on cold-water coral reefs. However, both the present results and previous observations indicate that *D. pertusum* skeletons harbour assemblages that differ from those of living coral and sediment (Yakimov et al., 2006; Van Bleijswijk et al., 2015). As observed in tropical reefs, these rubble-associated communities may contribute to biogeochemical processes and to microbial diversity at the reef scale (Beltrán et al., 2016; Sánchez-Quinto and Falcón, 2019; Wolfe et al., 2021).

Within *D. pertusum*, microbial communities differed significantly between branch and mucus samples and among colonies, although both factors explained relatively small proportions of the total variation (***Fig. 3B***). Colony identity accounted for more variation than compartment, and the absence of a significant interaction indicated that branch-mucus differences were consistent across colonies. This colony-level pattern was driven primarily by Colony B, whose microbial communities differed from both Colony A and Colony C, which did not differ significantly from each other. This pattern was also apparent in the relative abundance profiles of dominant families (***Fig. 3A***) and indicator taxa (***Fig. 4B***), and in the prevalence of core taxa (***Fig. 4A***). The data thus reveal a detectable but limited colony-level signal superimposed on substantial variation among samples within each colony.

All three colonies were sampled from the south-eastern side of Tisler Reef, where *D. pertusum* cover is higher than on the north-western side. This pattern is attributed to more favourable growth conditions associated with more frequent and prolonged downwelling, which transports warmer and chlorophyll-rich surface waters towards the reef (De Clippele et al., 2018). Tisler Reef is nonetheless a hydrodynamically complex environment in which current speed and direction vary over time and by position on the reef (De Clippele et al., 2018), and coral thickets can themselves modify near-bed flow, with potential consequences for particle encounter rates and coral feeding (Davies et al., 2009; Purser et al., 2010). Colonies A and C were separated by approximately 25 m horizontally and 4 m vertically, whereas Colony B was 40-57 m from the other colonies and 5-9 m deeper. Although small at the reef scale, these distances may still be sufficient to generate differences in flow exposure, food delivery, sedimentation, or local reef structure (Findlay et al., 2014; Corbera et al., 2022), offering one possible explanation for the differentiation of Colony B, although direct hydrodynamic and environmental measurements would be needed to test this hypothesis.

Similar biogeographic imprinting combined with weak host specificity has previously been reported for several Norwegian reefs (Schöttner et al., 2012). On Røst Reef specifically, the local similarity of coral-associated microbial assemblages from a same site was lost beyond the reef centre, with considerable variation in community structure towards the periphery (Schöttner et al., 2012). Mediterranean *D. pertusum* colonies have likewise shown extensive colony-to-colony microbiome variability within the same site and were thus hypothesized to be susceptible to environmental filtering (Meistertzheim et al., 2016). More direct evidence for environmental selection comes from Chapron et al. (2020), who found that bacterial communities of transplanted *D. pertusum* fragments shifted towards those of local colonies at the destination site. Together, these studies support the possibility that environmental differences operating over relatively small spatial scales contribute to among-colony microbiome variation.

Host-related factors may also contribute to the colony effect: host genotype has been shown to influence bacterial community composition in tropical corals, although its relative importance varies with environmental context (Glasl et al., 2019; Dubé et al., 2021). Such genotype effects could not be evaluated here but may be relevant, as neighbouring *D. pertusum* colonies are not necessarily genetically independent. Asexual propagation can produce spatially-extensive clones, and within the Skagerrak fjord system, including Tisler Reef, individual clones have been reported to extend over areas approaching 300 m^2^ (Dahl et al., 2012). However, the respective contributions of host relatedness and local environmental conditions in the present study cannot be separated without host genotyping and matched environmental measurements. The observed colony patterns should therefore be regarded as a basis for these hypotheses rather than as evidence for one specific underlying mechanism.

Within coral branches, both colony identity and position along the branch were associated with microbial community composition (***Fig. 3C-D***). Colony explained slightly more variation, but tip and bottom samples differed significantly from one another, while middle-position samples were not distinguishable from either group. Branch tips may experience greater exposure to ambient flow and particle delivery, whereas bottom sections are likely more sheltered within the colony structure (Purser et al., 2010). The sampled sections may also differ in branch orientation, polyp feeding activity, sediment exposure, tissue condition, or the relative contributions of tissue, mucus, and skeletal habitats (Sanna and Freiwald, 2021). These differences might be particularly pronounced for branch tips which harbour a polyp and thus a relatively important contribution of gastrovascular cavity tissues. In addition, tip and basal portions may differ in the age and physical characteristics of the underlying skeleton, potentially creating differences in the availability and physico-chemical characteristics of microbial micro-habitats (Ricci et al., 2019).

An early-identified feature of *D. pertusum* behaviour is the lack of co-ordination between polyps and of colony-wide communication (Shelton, 1980), suggesting that polyp-to-polyp differences may occur. As an azooxanthellate coral, *D. pertusum* relies strongly on heterotrophic feeding, and variation in food encounter and recent feeding history could plausibly contribute to local differences in its associated microbial assemblages. Distinctive bacterial communities have been detected within the gastrovascular cavities of tropical corals (Agostini et al., 2012; Bollati et al., 2024), and polyp-to-polyp microbiome variation has also been reported within Mediterranean *D. pertusum* colonies, which the authors attributed to variation in gastrovascular-cavity community due to differences in recently ingested food (Meistertzheim et al., 2016). Experimental evidence further showed that diet alters *D. pertusum* microbiome composition while retaining a species-specific pattern (Galand et al., 2020). Together, these findings offer a plausible explanation for fine-scale microbiome heterogeneity in *D. pertusum*, though the contribution of gastric contents to the microbial profiles obtained from whole-branch samples remains unknown. Sampling deep-sea coral colonies with ROV necessarily limits the precision with which coral branches can be sampled in a non-destructive manner and how microscale variables such as flow exposure, polyp density, sediment deposition, and orientation can be documented *in situ*. Despite this constraint, a significant difference between branch tip and bottom samples was detectable, suggesting that branch-scale spatial organisation warrants further investigation.

Microbial core and indicator species analyses further showed that prevalence within coral samples did not necessarily coincide with strict specificity to the coral habitat. Across the 42 identified coral core TFs, including the nine detected in 100% of coral branch samples (***Fig. 4A***), most were also highly prevalent and abundant in substrate, rubble, or seawater samples, with only five being more abundant in coral branches and two more abundant in coral mucus than in any other sample type (***Supplementary Table S4***). Therefore, they represent recurrent components of the branch-associated community, but not an environmentally exclusive coral core. Despite their widespread occurrence, these taxa might nevertheless have ecologically relevant interactions with the coral host, as microorganisms also detected in the surrounding environment can still be enriched within coral tissues and occupy functionally important niches (Ainsworth et al., 2015; Hernandez-Agreda et al., 2017; Van Oppen and Blackall, 2019). Indicator species analysis identified 15 TFs associated with branch samples, mucus samples, or both coral compartments relative to environmental samples (***Fig. 4B***). Specificity values were relatively high, with 73-98% of occurrences in the target groups, but fidelity was more moderate, with indicator taxa detected in 53-89% of target samples (***Supplementary Table S3***), suggesting these are preferential associations with coral samples without constituting universal microbiome members. Most also occurred at low relative abundance, which may have reduced their detection probability, although genuine biological patchiness among coral samples cannot be excluded. Only one TF was classified both as a core and indicator taxon. This limited overlap illustrates that prevalence on one hand, and specificity combined with fidelity on the other, capture different properties of the microbiome: a taxon can be widespread within coral samples without being coral-specific, or it can be preferentially associated with coral while occurring in only a subset of individuals.

### 4.2 Microbial taxa associated with *D. pertusum* reef environments

Taxonomic profiling of the dominant families at the reef and coral scale, complemented by the identification of putative core and indicator taxa, revealed potentially important microbial associates of this cold-water coral habitat.

Dominant families at the reef-scale fell into three broad patterns: seawater-enriched bacterioplankton families, benthic or substrate-associated groups, and families occurring widely across compartments (***Fig. 2A***). The bacterial taxa enriched in seawater, including Pelagibacterales SAR11 Clade I and Clade II (Alphaproteobacteria), Actinomarinaceae (Acidimicrobiia), and the SAR86 clade (Gammaproteobacteria), are characteristic members of globally distributed marine bacterioplankton, consistent with oligotrophic, oxygenated open water (Wietz et al., 2010; Giovannoni, 2017; López-Pérez et al., 2020; Roda-Garcia et al., 2023; Thomas et al., 2025). This indicates that seawater assemblages at Tisler Reef retained a taxonomic profile characteristic of open marine water columns, although the present study cannot quantify the potential contribution of reef-derived microorganisms to this signal.

Kiloniellaceae were prevalent across all reef habitats but dominated coral rubble communities, primarily represented by one uncultured taxon (15.16 ± 6.75% in relative abundance). The remaining Kiloniellaceae taxa showed contrasting distributions: one, identified as a *Thalassocola* sp., was prevalent in *D. pertusum* and coral rubble but absent from substrate and seawater; one, identified as *Tagaea marina*, was detected exclusively in coral branch samples; one, corresponding to *Kiloniella antarctica*, occurred only in seawater and coral mucus; and two, identified as *Limibacillus* sp. and *Pelagibius* sp., were found across all sample types. These contrasting distributions within a single family illustrate how family-level patterns can conceal taxa with distinct ecological niches. Woeseiaceae were similarly prevalent across all sample types, with increased abundance in coral rubble and substrate. Within this family, the dominant TF_6, identified as *Woeseia* sp., was prevalent across all sample types and was the top core taxon ubiquitously associated with *D. pertusum*, combining the rare attributes of being highly prevalent (100% in coral, substrate, and rubble samples) and abundant across many samples (5.9 ± 1.5% in rubble, 4.8 ± 1.0% in substrate, 1.6 ± 1.3% in coral branches, and 2.1 ± 1.7% in coral mucus, ***Supplementary Table S4***). Both Kiloniellaceae and Woeseiaceae have previously been reported as abundant members of marine sediments (Mußmann et al., 2017; Guerrero et al., 2022), notably in deep-sea areas where they have been hypothesised to play roles in nitrogen cycling (Modolon et al., 2025; Hageman et al., 2026). Although Kiloniellaceae and Woeseiaceae were the 6^th^ and 11^th^ most abundant families associated with *D. pertusum*, they have not previously been reported from other studies of this coral (***Table 1***).

Paracoccaceae was the most abundant bacterial family across all reef-scale compartments, represented by 158 taxa, and the second most abundant family in coral samples, represented by 134 taxa. This family has repeatedly been reported as a dominant *D. pertusum*-associated group (Rhodobacteraceae in older literature) (Neulinger et al., 2008; Kellogg et al., 2009, 2017; Chapron et al., 2021); we show, however, that it was widespread across all sample types, and especially abundant in seawater. This distribution suggests that Paracoccaceae may be better characterised as a globally abundant family of the reef environment than as an associate specifically selected by the coral host. Paracoccaceae have nonetheless been proposed to confer a nutritional advantage through mixotrophy and sulphur oxidation (Neulinger et al., 2008), but have also been flagged as an opportunistic, potentially dysbiosis-associated taxon whose relative abundance rises under thermal stress in *D. pertusum* (Chapron et al., 2021). Five of the identified core taxa belonged to Paracoccaceae, including *Sulfitobacter* (TF_94), *Limibaculum* (TF_39), and *Roseovarius* (TF_13), which were particularly abundant in coral samples compared to other samples (***Supplementary Table S4***). *Sulfitobacter* has previously been reported as an abundant microbial associate of *D. pertusum* and other deep-sea corals from a mid-shelf Norwegian reef (Jensen et al., 2019), whereas the other genera have not previously been reported from *D. pertusum* microbiomes (***Table 1***).

One of the most conspicuous taxonomic features of the *D. pertusum*-associated communities was the high relative abundance of Leptospiraceae (Spirochaetota) in a subset of branch and mucus samples. Of the four taxa within this family, three, identified as belonging to the genera *Leptospira* (detected only in coral branches), *Leptonema* (detected in branches and mucus), and *Turneriella* (detected in all sample types except seawater), occurred at low relative abundances, while the fourth and dominant taxon could only be resolved to family level. Although this dominant taxon was detected in all sample types, its relative abundance was much lower in substrate and rubble, supporting an association with the living coral compartments. Its distribution was nonetheless highly heterogeneous: Leptospiraceae occurred at low abundance in many coral samples, particularly in Colony B, but dominated others, particularly in tip branches from Colony C (***Supplementary Figure S2***). Its prominence therefore reflected strong but spatially patchy dominance rather than uniformly high abundance across colonies or within colonies.

Leptospiraceae has not previously been directly reported in *D. pertusum* studies (***Table 1***), though the family is not entirely unprecedented in coral- and deep-sea-associated systems: a Leptospiraceae TF from the red coral *Corallium rubrum* clustered with clones retrieved from *D. pertusum* and *Montastraea faveolata*, and with cold-water environmental sequences (Sunagawa et al., 2009; Van De Water et al., 2016). *Leptospira* has also been identified as a stable microbial associate of gorgonians in the Mediterranean Sea (van de Water et al., 2018), and Leptospiraceae have been listed among abundant families associated with the coral *Pocillopora acuta* (Epstein et al., 2019), with early life stages of *Acropora tenuis* (Damjanovic et al., 2020), and as specific bacterial associates of a temperate sponge species (Lamb and Watts, 2023). Beyond these reports, Leptospiraceae remain largely unknown as coral associates, and their potential roles on cold-water reefs remain unclear. They have nevertheless been detected in various marine environmental samples, with evidence suggesting a role in ß-oxidation of fatty acids, notably of oleic acid, a major fatty acid of marine phytoplankton (Wakeham et al., 1997; Yu et al., 2023).

Despite the variability in dominant taxa reported from the literature on *D. pertusum* (summarized in ***Table 1***), and the within-colony variability of taxa identified through our study, some associations across independent studies and geographic regions point to several bacterial lineages that may be consistently linked to *D. pertusum* across its range. Several other families and genera identified in the present study also recurred among prior reports of dominant *D. pertusum*-associated taxa (***Table 1***). *Pseudahrensias* sp., identified here as a core taxon within Rhizobiaceae, the third most abundant family associated with the coral, has previously been reported as highly dominant in dead *D. pertusum* skeletons (Van Bleijswijk et al., 2015). *Colwellia* sp. (TF_2), the second most abundant TF across all coral samples and a member of the fourth most abundant family, Colwelliaceae, has rarely been reported from *D. pertusum*, with the exception of a bacterial isolate from coral tissue, skeleton, and mucus homogenates corals in the northern Gulf of Mexico (Galkiewicz et al., 2011). The class Bacteroidia, which includes Flavobacteriaceae, the fifth most abundant family here, has previously been reported as dominant in *D. pertusum* mucus and tissues from several Atlantic locations (Van Bleijswijk et al., 2015; Chemel et al., 2024; Zanotti et al., 2024). Alteromonadaceae and Vibrionaceae, both particularly enriched in Colony B, have previously been observed to increase in relative abundance in heat-stressed *D. pertusum* (Chemel et al., 2024), although Vibrionaceae have also been reported to dominate in healthy corals and have been hypothesised to play beneficial roles in host immunity and nutrition (Galkiewicz et al., 2011).

## 5 Conclusion

Taken together, these results indicate that *D. pertusum* at Tisler Reef harbours microbial assemblages that are distinguishable from those in the surrounding seawater and benthic habitats, but that do not comprise a strong microbial signature with an environmentally exclusive taxonomic core. Instead, the coral-associated microbiome combines recurrent, environmentally widespread taxa with less prevalent taxa that show stronger associations with coral branches or mucus. Its composition also varies among neighbouring colonies and within individual colonies, suggesting that coral-associated communities are structured but responsive to variation occurring at relatively fine spatial and biological scales. These patterns are consistent with a microbiome that is structured, yet potentially responsive to host state and local environmental conditions, rather than one that is taxonomically invariant across the coral population. Such flexibility could help *D. pertusum* respond rapidly to changing environmental conditions (Voolstra and Ziegler, 2020), and may also reflect sensitivity to micro-scale factors such as current exposure, sediment deposition, and nutritional status of individual polyps (Galand et al., 2020). Identifying the drivers of the fine-scale variability will require sampling designs that pair microbiome characterisation with host genotyping and direct environmental measurements. More broadly, these findings underscore that the microbiome of the habitat-forming cold-water coral *D. pertusum* cannot be reduced to a single, well-defined, and consistent community. It is rather best understood as a variable assemblage likely shaped by finer-scale conditions experienced by individual colonies and even individual sections within a single colony. Uncovering what factors influence cold-water coral microbiomes and characterising this variability, rather than searching solely for a fixed core, will be increasingly important to understand how these corals function and how they might respond to the environmental pressures they face.

## 6 Conflict of Interest

The authors declare that the research was conducted in the absence of any commercial or financial relationships that could be construed as a potential conflict of interest.

## 7 Author Contributions

PF and CP conceived and designed the experiments with the contributions of AL. Data was collected by CP with the contributions of AIL, SMS, NEV, and SD. CS analysed the data and wrote the paper. PF, AL, NEV, EH, and MK contributed to manuscript revision.

## 8 Funding

This research was funded through the 2020-2021 BiodivERsA COFUND call for research proposals, with the national funders FWF - Austrian Science Fund (project number I 5838-B) and Swedish Environmental Protection Agency (2022-00025).

## 9 Acknowledgments

This work was supported by the European Union project RESTORESEAS (EU-BiodivERsA BiodivRestore-253), which is funded through the BiodivRestore joint call, a collaborative initiative between Biodiversa + and the Water Joint Programming Initiative (JPI). We thank Alexandra Wanka and Sandra Kirchner (Central Research Laboratories of NHMW) for support during laboratorial work.

## 10 Data Availability Statement

The raw 16S rRNA gene amplicon sequencing data have been deposited in the NCBI Sequence Read Archive (SRA) database under BioProject accession PRJNA1543492.

Metadata, taxonomic feature, and statistical data and are available via the Zenodo repository: https://doi.org/10.5281/zenodo.23239695

## Notes

https://doi.org/10.5281/zenodo.23239695

### Competing Interest Statement

The authors have declared no competing interest.

