## Supplementary Material for "High variability and limited host specificity characterise the microbiome of *Desmophyllum pertusum* across a cold-water coral reef"

### 1 Supplementary Material & Methods S1

**Table S1.** Number of samples retained for 16S microbial community analysis across the three sites and the different sample types.

| Sample Type | | | Colony A | | Colony B | | Colony C | | TOTAL | |
|---|---|---|---|---|---|---|---|---|---|---|
| Healthy Coral | Branches | Tip | 6 | total: 16 | 4 | total: 14 | 7 | total: 17 | **17** | total: **47** |
| | | Middle | 6 | | 5 | | 5 | | **16** | |
| | | Bottom | 4 | | 5 | | 5 | | **14** | |
| | Mucus | | 7 | | 5 | | 5 | | **17** | |
| Coral Rubble | Branches | | 3 | | 4 | | 4 | | **11** | |
| Environment | Substrate | Sediment | 2 | | – | | – | | **7** | |
| | | Rock / Shell | – | | 2 | | 3 | | | |
| | Seawater | | 3 | | 3 | | 3 | | **9** | |
| | | **TOTAL** | **31** | | **28** | | **32** | | **91** | |

### 2 Supplementary Data S2

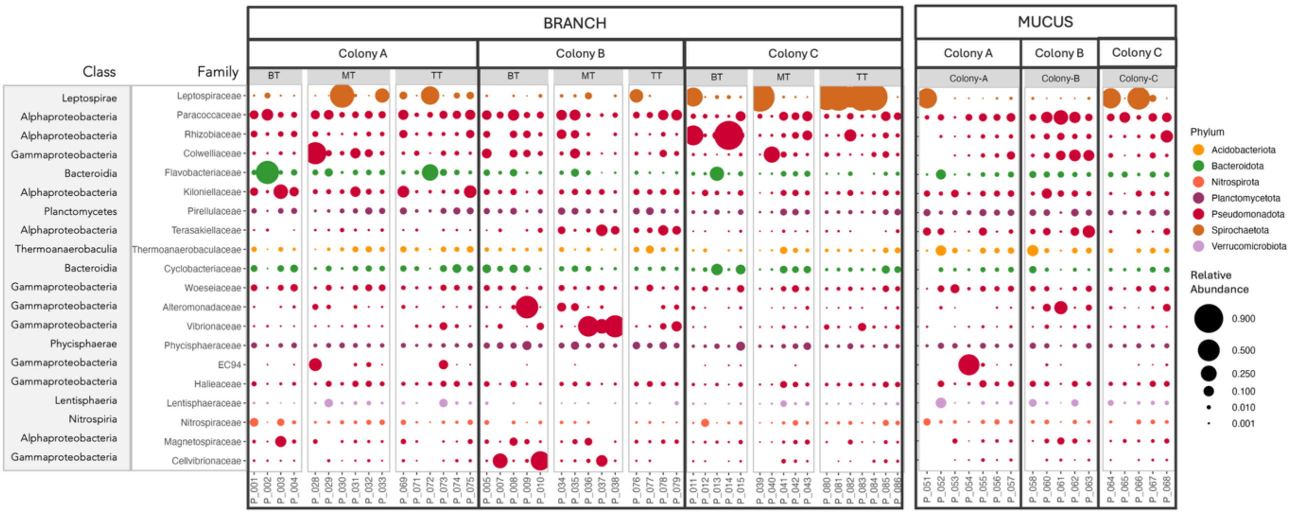


**Fig. S2.** Relative abundance of the 20 most abundant prokaryotic families across all coral samples. Samples are grouped into branch and mucus samples and further divided in colony groups. Coral branch samples are grouped according to the position on the branch: BT, bottom of the branch; MT, middle of the branch; TT, tip of the branch. Point size represents the relative abundance of each family in each sample, and the colours indicate the phyla of the family.

### 3 Supplementary Data S3 – Indicator Species Analysis Statistics

**Table S3. Indicator species analysis statistics.** Report of the results from the indicator species analysis using the *multipatt* function with 999 permutations (*indicspecies* R package). Indicator taxa were filtered based on the threshold p-value ≤ 0.01 with minimum specificity (component A) and minimum fidelity (component B) set to 50%.

| TF_Rank | Taxa | Indicator Group | stat | p.value | A | B |
|---|---|---|---|---|---|---|
| TF_145 | Neorickettsia_s | Branch | 0.7457 | 0.001 | 90.1% | 61.7% |
| TF_379 | Persicirhabdus_s | Mucus | 0.6945 | 0.001 | 82.0% | 58.8% |
| TF_280 | Rubritalea_s | Mucus | 0.7760 | 0.001 | 73.1% | 82.4% |
| TF_349 | Pseudomonas putida | Mucus | 0.7828 | 0.001 | 86.8% | 70.6% |
| TF_223 | Kangiella chungangensis | Mucus | 0.6755 | 0.006 | 70.5% | 64.7% |
| TF_309 | Winogradskyella_s | Mucus | 0.7363 | 0.001 | 76.8% | 70.6% |
| TF_118 | Lentimonas_s | Branch+Mucus | 0.8227 | 0.01 | 86.6% | 78.1% |
| TF_54 | Porticoccus_s | Branch+Mucus | 0.8614 | 0.005 | 89.6% | 82.8% |
| TF_247 | Pelagicola_s | Branch+Mucus | 0.7655 | 0.002 | 96.2% | 60.9% |
| TF_204 | Sinobacterium_s | Branch+Mucus | 0.7207 | 0.003 | 97.8% | 53.1% |
| TF_187 | Hyphomonadaceae_g_s | Branch+Mucus | 0.7294 | 0.006 | 94.6% | 56.3% |
| TF_14 | Marivibrio_s | Branch+Mucus | 0.9187 | 0.001 | 94.8% | 89.1% |
| TF_87 | FS140-16B-02 marine group_s | Branch+Mucus | 0.8631 | 0.003 | 86.7% | 85.9% |
| TF_53 | Phycisphaera_s | Branch+Mucus | 0.8711 | 0.004 | 93.4% | 81.3% |
| TF_150 | JL-ETNP-F27_s | Branch+Mucus | 0.7937 | 0.005 | 87.6% | 71.9% |

### 4 Supplementary Data S4 – Indicator and Core Taxa Relative Abundances and Prevalence

**Table S4. Relative abundance and prevalence of core and indicator microbial taxa associated with *D. pertusum*.** Taxa are defined by TF number, corresponding to their rank in relative abundance across all coral samples, followed by their lowest available taxonomic classification. Unresolved taxonomic ranks are indicated by suffixes: "_c" (class), "_o" (order), "_f" (family), "_g" (genus), and "_s" (species). The "Core" and "Indic" columns indicate if taxa were identified as core taxa and / or indicator taxa. Asterix in the "Core" column denote the nine taxa present in 100% of coral branch samples across all three colonies. Column abbreviations designate the different sample types: "Bra" (coral branch), "Muc" (coral mucus), "Rub" (coral rubble), "Sub" (substrate), "Wat" (water). Relative abundance and prevalence values are expressed as percentages (%).

| | | | | Relative Abundance (%) | | | | | | | | | | Prevalence (%) | | | | |
|---|---|---|---|---|---|---|---|---|---|---|---|---|---|---|---|---|---|---|
| | | | | Mean | | | | | Stdev | | | | | | | | | |
| TF_Rank | Taxa | Core | Indic | Bra | Muc | Rub | Sub | Wat | Bra | Muc | Rub | Sub | Wat | Bra | Muc | Rub | Sub | Wat |
| **TF_6** | Woeseia _s | Yes* | | 1.63 | 2.06 | 5.93 | 4.79 | 0.37 | 1.27 | 1.68 | 1.49 | 1.01 | 0.35 | 100.0 | 100.0 | 100.0 | 100.0 | 77.8 |
| **TF_8** | NB1-j _c_o_f_g_s | Yes* | | 1.62 | 1.76 | 4.16 | 3.47 | 0.24 | 1.24 | 1.16 | 1.13 | 1.50 | 0.31 | 100.0 | 100.0 | 100.0 | 100.0 | 77.8 |
| **TF_21** | OM190 _o_f_g_s | Yes* | | 0.96 | 1.05 | 1.21 | 2.28 | 0.44 | 0.99 | 0.76 | 0.34 | 0.64 | 0.35 | 100.0 | 94.1 | 100.0 | 100.0 | 77.8 |
| **TF_25** | Gammaproteobacteria _o_f_g_s | Yes* | | 0.80 | 0.83 | 0.94 | 1.63 | 0.18 | 0.72 | 0.56 | 0.36 | 0.76 | 0.79 | 100.0 | 100.0 | 100.0 | 100.0 | 66.7 |

| | | | | | | | | | | | | | | | | | |
|---|---|---|---|---|---|---|---|---|---|---|---|---|---|---|---|---|---|
| **TF_26** | Rubinisphaeraceae _g_s | Yes* | | 0.64 | 1.02 | 0.14 | 0.60 | 1.11 | 0.98 | 1.00 | 0.08 | 0.44 | 2.97 | 100.0 | 100.0 | 100.0 | 100.0 | 88.9 |
| **TF_27** | Paracoccaceae _g_s | Yes* | | 0.63 | 1.02 | 0.68 | 0.41 | 3.41 | 0.57 | 0.95 | 1.23 | 0.15 | 0.24 | 100.0 | 94.1 | 100.0 | 100.0 | 100.0 |
| **TF_31** | Pirellulaceae _g_s | Yes* | | 0.65 | 0.76 | 0.65 | 1.42 | 0.23 | 0.51 | 0.49 | 0.23 | 0.28 | 0.32 | 100.0 | 94.1 | 100.0 | 100.0 | 77.8 |
| **TF_32** | OM60(NOR5) clade _s | Yes* | | 0.59 | 0.88 | 0.20 | 0.71 | 0.43 | 0.49 | 0.65 | 0.10 | 0.28 | 9.26 | 100.0 | 94.1 | 100.0 | 100.0 | 77.8 |
| **TF_47** | Clade Ia _s | Yes* | | 0.25 | 0.92 | 0.19 | 0.09 | 11.54 | 0.34 | 2.49 | 0.17 | 0.06 | 3.90 | 100.0 | 94.1 | 100.0 | 100.0 | 88.9 |
| **TF_2** | Colwellia _s | Yes | | 2.87 | 2.66 | 0.37 | 3.40 | 1.40 | 6.67 | 3.92 | 0.30 | 6.19 | 0.53 | 97.9 | 88.2 | 100.0 | 100.0 | 77.8 |
| **TF_3** | UBA10353 marine group _f_g_s | Yes | | 2.40 | 3.94 | 3.15 | 0.92 | 0.82 | 8.72 | 9.24 | 5.66 | 0.63 | 0.12 | 95.7 | 88.2 | 100.0 | 100.0 | 88.9 |
| **TF_4** | Pseudahrensia _s | Yes | | 3.12 | 0.28 | 0.65 | 0.87 | 0.07 | 14.21 | 0.33 | 0.77 | 0.39 | 0.52 | 91.5 | 82.4 | 100.0 | 100.0 | 55.6 |
| **TF_7** | Rhizobiaceae _g_s | Yes | | 1.78 | 1.55 | 0.18 | 0.30 | 0.27 | 2.71 | 3.20 | 0.14 | 0.25 | 0.77 | 95.7 | 94.1 | 100.0 | 100.0 | 66.7 |
| **TF_10** | Kiloniellaceae _g_s | Yes | | 1.71 | 1.28 | 15.16 | 2.83 | 0.41 | 2.89 | 1.16 | 6.75 | 2.13 | 0.63 | 97.9 | 100.0 | 100.0 | 100.0 | 77.8 |
| **TF_11** | Cyclobacteriaceae _g_s | Yes | | 1.79 | 0.70 | 2.23 | 1.40 | 0.41 | 2.02 | 0.61 | 0.85 | 0.47 | 0.45 | 97.9 | 88.2 | 100.0 | 100.0 | 77.8 |
| **TF_12** | Subgroup 10 _s | Yes | | 1.07 | 2.53 | 1.45 | 1.80 | 0.19 | 1.04 | 3.19 | 0.40 | 0.32 | 0.12 | 97.9 | 94.1 | 100.0 | 100.0 | 66.7 |
| **TF_13** | Roseovarius _s | Yes | | 1.17 | 2.14 | 0.98 | 0.13 | 0.06 | 1.69 | 3.48 | 2.85 | 0.07 | 0.72 | 91.5 | 82.4 | 100.0 | 100.0 | 55.6 |
| **TF_17** | Unknown Family _g_s | Yes | | 1.04 | 1.56 | 0.66 | 1.92 | 0.34 | 1.00 | 1.04 | 0.27 | 0.55 | 0.25 | 97.9 | 94.1 | 100.0 | 100.0 | 66.7 |
| **TF_18** | Haliangiaceae _g_s | Yes | | 1.50 | 0.27 | 0.96 | 1.11 | 0.14 | 2.95 | 0.33 | 0.44 | 0.94 | 0.17 | 93.6 | 82.4 | 100.0 | 100.0 | 66.7 |
| **TF_20** | Pelagibius _s | Yes | | 1.04 | 0.96 | 0.89 | 0.44 | 0.07 | 2.20 | 1.77 | 0.24 | 0.15 | 0.37 | 93.6 | 88.2 | 100.0 | 100.0 | 55.6 |
| **TF_22** | Nitrospira _s | Yes | | 0.93 | 1.06 | 4.11 | 1.94 | 0.17 | 1.34 | 0.98 | 1.63 | 0.44 | 0.23 | 93.6 | 94.1 | 100.0 | 100.0 | 77.8 |
| **TF_30** | Nitrosomonas _s | Yes | | 0.74 | 0.52 | 1.67 | 1.04 | 0.31 | 0.77 | 0.38 | 0.67 | 0.30 | 0.17 | 97.9 | 94.1 | 100.0 | 100.0 | 77.8 |
| **TF_34** | Anderseniella _s | Yes | | 0.60 | 0.47 | 1.34 | 1.12 | 0.07 | 0.87 | 0.37 | 0.53 | 0.49 | 1.16 | 93.6 | 88.2 | 100.0 | 100.0 | 55.6 |
| **TF_36** | Defluviicoccales _f_g_s | Yes | | 0.51 | 0.68 | 2.36 | 1.29 | 1.16 | 0.80 | 0.89 | 1.31 | 1.06 | 0.33 | 93.6 | 100.0 | 100.0 | 100.0 | 77.8 |
| **TF_37** | Roseobacter clade NAC11-7 lineage _s | Yes | | 0.46 | 0.69 | 0.53 | 0.18 | 0.18 | 0.56 | 0.88 | 1.43 | 0.09 | 0.21 | 91.5 | 88.2 | 100.0 | 100.0 | 66.7 |
| **TF_38** | Pir4 lineage _s | Yes | | 0.49 | 0.59 | 0.46 | 0.79 | 0.10 | 0.47 | 0.45 | 0.19 | 0.32 | 0.16 | 93.6 | 88.2 | 100.0 | 100.0 | 66.7 |
| **TF_39** | Limibaculum _s | Yes | | 0.51 | 0.51 | 0.24 | 0.77 | 0.06 | 0.58 | 0.52 | 0.11 | 0.29 | 0.31 | 91.5 | 94.1 | 100.0 | 100.0 | 55.6 |
| **TF_40** | Sandaracinaceae _g_s | Yes | | 0.44 | 0.67 | 0.26 | 0.73 | 0.16 | 0.51 | 0.47 | 0.11 | 0.24 | 0.08 | 95.7 | 100.0 | 100.0 | 100.0 | 66.7 |
| **TF_41** | Kangiellaceae _g_s | Yes | | 0.56 | 0.28 | 0.18 | 0.48 | 0.03 | 0.83 | 0.29 | 0.16 | 0.35 | 0.25 | 91.5 | 82.4 | 100.0 | 100.0 | 66.7 |
| **TF_49** | Coxiella _s | Yes | | 0.40 | 0.48 | 0.25 | 0.41 | 0.29 | 0.55 | 0.67 | 0.10 | 0.14 | 0.05 | 91.5 | 82.4 | 100.0 | 100.0 | 77.8 |
| **TF_50** | Subgroup 22 _o_f_g_s | Yes | | 0.41 | 0.44 | 0.57 | 1.01 | 0.03 | 0.58 | 0.56 | 0.21 | 0.35 | 0.47 | 95.7 | 88.2 | 100.0 | 100.0 | 55.6 |
| **TF_52** | BD2-11 terrestrial group _o_f_g_s | Yes | | 0.40 | 0.45 | 0.80 | 1.06 | 0.57 | 0.51 | 0.59 | 0.33 | 0.46 | 0.50 | 93.6 | 94.1 | 100.0 | 100.0 | 88.9 |
| **TF_54** | Saprospiraceae _g_s | Yes | Yes | 0.41 | 0.36 | 0.27 | 1.84 | 0.27 | 0.45 | 0.41 | 0.10 | 1.12 | 0.12 | 95.7 | 94.1 | 100.0 | 100.0 | 77.8 |
| **TF_58** | Sva0081 sediment group _s | Yes | | 0.32 | 0.57 | 0.12 | 0.36 | 0.08 | 0.33 | 0.66 | 0.07 | 0.09 | 0.20 | 93.6 | 94.1 | 100.0 | 100.0 | 66.7 |
| **TF_59** | Mariniblastus _s | Yes | | 0.31 | 0.56 | 0.46 | 0.51 | 0.32 | 0.34 | 0.56 | 0.62 | 0.15 | 0.80 | 95.7 | 100.0 | 100.0 | 100.0 | 88.9 |
| **TF_63** | OM43 clade _s | Yes | | 0.36 | 0.34 | 0.07 | 0.07 | 0.84 | 0.40 | 0.51 | 0.10 | 0.05 | 0.71 | 91.5 | 88.2 | 90.9 | 85.7 | 77.8 |
| **TF_64** | Gimesiaceae _g_s | Yes | | 0.23 | 0.60 | 0.19 | 0.43 | 0.77 | 0.19 | 0.63 | 0.14 | 0.14 | 0.08 | 97.9 | 94.1 | 100.0 | 100.0 | 77.8 |
| **TF_72** | Seongchinamella _s | Yes | | 0.26 | 0.42 | 0.10 | 0.40 | 0.06 | 0.27 | 0.41 | 0.05 | 0.12 | 0.09 | 97.9 | 94.1 | 100.0 | 100.0 | 66.7 |
| **TF_74** | Arenicellaceae _g_s | Yes | | 0.27 | 0.34 | 0.59 | 0.74 | 0.12 | 0.26 | 0.26 | 0.28 | 0.25 | 0.14 | 93.6 | 94.1 | 100.0 | 100.0 | 77.8 |
| **TF_92** | Legionellaceae _g_s | Yes | | 0.21 | 0.30 | 0.04 | 0.09 | 0.13 | 0.23 | 0.39 | 0.05 | 0.05 | 1.13 | 91.5 | 82.4 | 72.7 | 100.0 | 66.7 |
| **TF_94** | Sulfitobacter _s | Yes | | 0.26 | 0.15 | 0.11 | 0.04 | 0.46 | 0.66 | 0.18 | 0.26 | 0.03 | 3.97 | 93.6 | 82.4 | 54.5 | 85.7 | 77.8 |
| **TF_117** | SAR86 clade _g_s | Yes | | 0.16 | 0.16 | 0.21 | 0.07 | 4.06 | 0.28 | 0.50 | 0.50 | 0.05 | 0.06 | 95.7 | 88.2 | 100.0 | 100.0 | 88.9 |
| **TF_14** | Marivibrio _s | | Yes | 1.91 | 0.01 | 0.01 | 0.00 | 0.02 | 8.32 | 0.03 | 0.03 | 0.00 | 0.07 | 23.4 | 11.8 | 9.1 | 0.0 | 11.1 |
| **TF_53** | Phycisphaera _s | | Yes | 0.42 | 0.38 | 0.13 | 0.21 | 0.03 | 0.65 | 0.68 | 0.07 | 0.12 | 0.02 | 85.1 | 70.6 | 100.0 | 100.0 | 44.4 |
| **TF_87** | FS140-16B-02 marine group _s | | Yes | 0.21 | 0.32 | 0.15 | 0.33 | 0.02 | 0.25 | 0.31 | 0.13 | 0.15 | 0.11 | 85.1 | 88.2 | 100.0 | 100.0 | 55.6 |
| **TF_118** | Lentimonas _s | | Yes | 0.15 | 0.19 | 0.05 | 0.17 | 0.05 | 0.20 | 0.21 | 0.04 | 0.06 | 0.09 | 68.1 | 70.6 | 81.8 | 100.0 | 55.6 |
| **TF_145** | Neorickettsia _s | | Yes | 0.11 | 0.17 | 0.06 | 0.15 | 0.11 | 0.15 | 0.23 | 0.04 | 0.08 | 0.21 | 59.6 | 82.4 | 90.9 | 100.0 | 88.9 |
| **TF_150** | JL-ETNP-F27 _s | | Yes | 0.12 | 0.11 | 0.01 | 0.08 | 0.07 | 0.33 | 0.18 | 0.01 | 0.09 | 0.06 | 48.9 | 58.8 | 27.3 | 71.4 | 33.3 |
| **TF_187** | Hyphomonadaceae _g_s | | Yes | 0.06 | 0.17 | 0.48 | 0.32 | 0.02 | 0.10 | 0.53 | 0.15 | 0.14 | 0.00 | 59.6 | 58.8 | 100.0 | 100.0 | 11.1 |
| **TF_204** | Sinobacterium _s | | Yes | 0.08 | 0.08 | 0.27 | 0.21 | 0.00 | 0.11 | 0.12 | 0.12 | 0.06 | 0.20 | 55.3 | 64.7 | 100.0 | 100.0 | 11.1 |
| **TF_223** | Kangiella chungangensis | | Yes | 0.08 | 0.02 | 0.01 | 0.03 | 0.18 | 0.27 | 0.02 | 0.02 | 0.01 | 0.14 | 59.6 | 58.8 | 36.4 | 100.0 | 77.8 |
| **TF_247** | Pelagicola _s | | Yes | 0.05 | 0.07 | 0.00 | 0.03 | 0.06 | 0.08 | 0.13 | 0.00 | 0.03 | 0.00 | 44.7 | 41.2 | 0.0 | 57.1 | 44.4 |
| **TF_280** | Rubritalea _s | | Yes | 0.04 | 0.04 | 0.00 | 0.00 | 0.00 | 0.26 | 0.16 | 0.00 | 0.00 | 0.01 | 8.5 | 17.6 | 0.0 | 0.0 | 0.0 |
| **TF_309** | Winogradskyella _s | | Yes | 0.02 | 0.04 | 0.09 | 0.06 | 0.00 | 0.05 | 0.07 | 0.05 | 0.02 | 0.22 | 38.3 | 52.9 | 100.0 | 100.0 | 11.1 |
| **TF_349** | Pseudomonas putida | | Yes | 0.03 | 0.01 | 0.00 | 0.12 | 0.07 | 0.18 | 0.02 | 0.00 | 0.31 | 0.00 | 6.4 | 17.6 | 0.0 | 14.3 | 11.1 |
| **TF_379** | Persicirhabdus _s | | Yes | 0.02 | 0.01 | 0.00 | 0.00 | 0.00 | 0.05 | 0.03 | 0.01 | 0.01 | 0.00 | 23.4 | 23.5 | 18.2 | 28.6 | 0.0 |
